# Distance-Restraint-Guided Diffusion Models for Sampling Protein Conformational Changes and Ligand Dissociation Pathways

**DOI:** 10.64898/2026.01.30.702714

**Authors:** Tatsuki Hori, Yoshitaka Moriwaki, Ryuichiro Ishitani

**Author notes:** To whom reprint requests should be addressed.

## Abstract

Protein conformational dynamics and ligand binding processes are fundamental to biological function, yet their systematic sampling and thermodynamic characterization remain challenging. Here, we present a distance-restraint-guided inference method that extends AlphaFold3-like diffusion model frameworks to predict protein structures at specified conformational states. By restraining intergroup distances defined as center-of-mass distances between atom groups during the reverse diffusion process, our method enables systematic sampling along reaction coordinates without model retraining. We implemented this approach in Boltz-2 and demonstrated its effectiveness on three model proteins that undergo open-closed conformational transitions, as well as on a protein-peptide dissociation pathway. Compared with conventional approaches that induce conformational diversity by manipulating input multiple sequence alignments, our method achieved more uniform coverage of conformational space while maintaining high structural quality as assessed by both learning-based confidence metrics and stereochemistry-based validation. By combining distance-restrained sampling with molecular dynamics simulations, we constructed free energy landscapes and quantitatively estimated binding free energies. Altogether, our approach bridges deep learning-based structure prediction and physics-based simulations, providing an efficient strategy for characterizing the dynamics of biomolecules. The materials supporting this article, including the source code and Colab notebook are available at github (https://github.com/cddlab/boltz_restr).

## Introduction

Protein structures are not static entities but exhibit anisotropic and anharmonic conformational changes, and these dynamic behaviors are intimately linked to biological function^1–3^. Elucidating the structural transitions of proteins and their underlying free energy landscapes is indispensable for understanding enzymatic reaction mechanisms, allosteric regulation, and structure-based drug design^4,5^. Various approaches based on molecular dynamics (MD) simulations have been developed to explore the conformational changes of biomacromolecules. Classical MD simulations can generate physicochemically consistent structural ensembles by tracking atomic trajectories according to Newtonian equations of motion. However, many biologically relevant conformational changes occur on timescales ranging from microseconds to milliseconds, making sufficient sampling difficult with conventional MD simulations due to computational cost constraints. To address this challenge, numerous enhanced sampling methods have been proposed, including Gaussian accelerated MD (GaMD)^6^, metadynamics^7^, and Parallel Cascade Selection MD (PaCS-MD)^8^. These methods enable efficient exploration of physically plausible conformational transition pathways. Combined with statistical mechanical treatments such as Markov state models (MSM)^9^ and the multistate Bennett acceptance ratio (MBAR) method^10^, they allow quantitative evaluation of free energy changes associated with structural transitions. Nevertheless, exhaustive conformational sampling in complex biomolecular systems still requires enormous computational resources.

Since the release of AlphaFold2^11^, methods for predicting conformational changes using deep learning-based structure prediction have rapidly advanced. In particular, approaches that control coevolutionary signals by manipulating the input Multiple Sequence Alignment (MSA) to generate different conformational states (MSA modulation methods) have attracted considerable attention. Several strategies have been proposed, including random subsampling of MSAs^12^, alanine substitution of residues thought to be involved in conformational changes^13^, MSA clustering^14^, and random masking of amino acid residues^15^. These methods achieve conformational sampling that is remarkably faster than conventional simulation approaches. They have successfully generated multiple functionally important conformational states, such as the open and closed states of membrane transporters^12^. This is consistent with the hypothesis that MSA-encoded coevolutionary information guides AlphaFold2/3 to search globally for low-energy structures within the vast conformational energy landscape^16^.

Recent advances in generative models have opened new perspectives for conformational sampling of biomolecules^17,18^. There is a useful conceptual parallel between generative models and statistical mechanics. Just as image generation models sample from learned probability distributions over images, MD simulations sample from the Boltzmann distribution of biomolecular systems through physics-based simulations governed by the Newtonian equations of motion. From this perspective, if deep learning-based generative models can learn the Boltzmann distribution of biomolecular systems, structural ensembles can be efficiently sampled through one-shot generation without performing physical simulations. Furthermore, free energy calculations could also become possible by directly evaluating the likelihood of the generative distribution. Several pioneering studies have been reported in this direction. Boltzmann generators employ invertible neural networks to generate statistically independent samples from the Boltzmann distribution in a one-shot manner^17^. AlphaFold3 and its clones generate atomic coordinates using diffusion models, enabling prediction of complex structures containing proteins and small molecules^19^. These models learn the distribution of their training data, which comprises static protein structures. Consequently, they are unlikely to capture the Boltzmann distribution of dynamic biomolecules at ambient temperature and pressure with sufficient accuracy to quantitatively predict free energy changes. Another diffusion model-based approach is BioEmu (Biomolecular Emulator), which generates protein structures reflecting thermodynamic states by learning from experimental structures and MD simulation trajectories with thermodynamic weighting^18^. However, BioEmu outputs only backbone structures, requiring separate calculations for side-chain conformations, and cannot handle non-protein compounds or ligands. Given that current generative models either lack thermodynamic accuracy or have limited applicability to complex biomolecular systems, combining physics-based simulation methods with deep learning-based structure prediction provides a practical solution for quantitative evaluation of conformational changes.

The diffusion model in AlphaFold3 allows the incorporation of arbitrary restraints on atomic coordinates at each step of the reverse diffusion process during inference^20^. Using this restraint-guided inference method, previous work demonstrated that stereochemical errors in ligands can be effectively suppressed by conformer restraints during reverse diffusion steps. In this study, we extended this framework by proposing a distance-restraint-guided inference method (hereafter, distance-restraint method) that predicts structures corresponding to specified states by restraining intergroup distances to target values during the reverse diffusion process. By combining this method with MD simulations, we developed an approach that enables quantitative estimation of free energy profiles along reaction coordinates. To validate the versatility of this strategy, we employed two distinct thermodynamic analysis frameworks: MSM^9^ and umbrella sampling (US) with MBAR method^10,21^. We demonstrated the effectiveness of this method by applying it to protein conformational changes and ligand dissociation processes.

## Methods

### Implementation of the distance-based restraints

The three-dimensional structure of proteins changes in response to both intramolecular interactions between domains and intermolecular interactions with other biomolecules such as ligands. In this study, we employed the intergroup distance *d*, defined as the center-of-mass distance between a pair of atom groups, as a reaction coordinate to characterize these structural changes. Based on this framework, we developed a method to predict three-dimensional structures at specified values of *d*. For simplicity, we focus on a single atom group pair with distance *d*, although the theoretical framework is readily extensible to multidimensional reaction coordinates involving multiple intergroup distances (Figure 1).

**Figure 1.**
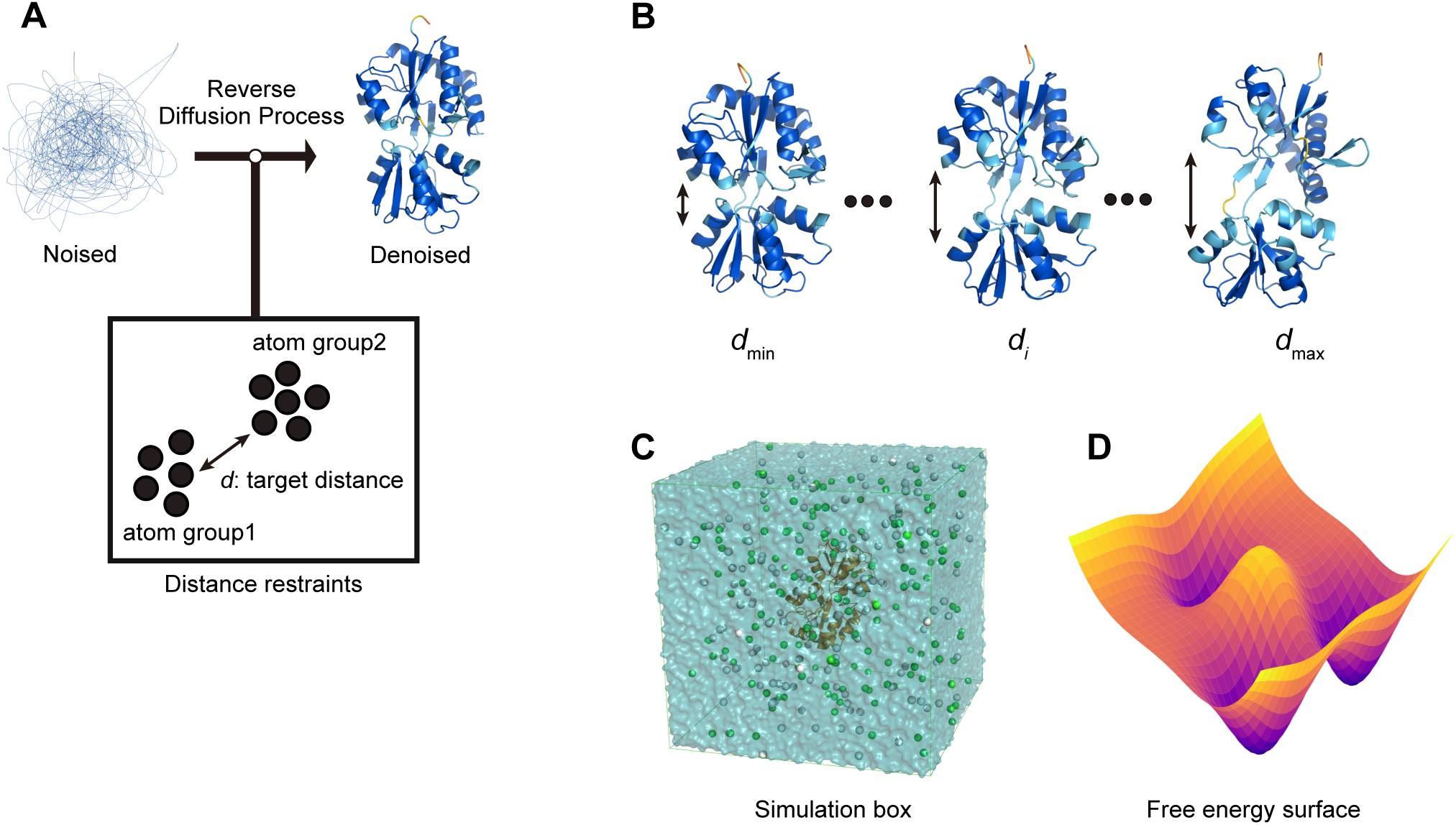
Overview of the distance-restraint-guided inference method and free energy analysis workflow. (A) Schematic of the distance-restraint method. During the reverse diffusion process, intergroup distances between defined atom groups are restrained to target values, directing the generation toward specified conformational states. (B) Systematic sampling of protein conformational states by varying the target distance from *d*_min_ to *d*_max_. Structures at intermediate distances (*d_i_*) are also generated to cover the conformational transition pathway. (C) Representative MD simulation system showing a protein structure solvated in an explicit water box with ions. (D) Schematic of a two-dimensional free energy landscape calculated from the results of the MD simulations.

The AlphaFold3-like structure prediction framework underlying our method uses a diffusion model to generate atomic coordinates. During inference, this model performs a reverse diffusion process that progressively refines atomic coordinates from noise. Our previous work demonstrated that predicted structures can be controlled by introducing chemical restraints such as bond lengths, bond angles, and chiral volumes during the reverse diffusion process^20^. In the present study, we extended this restraint-guided inference framework to predict structures in desired conformational states by restraining the intergroup distance *d* to a target value during the reverse diffusion process.

We defined two distinct atom groups {*a*_1_} and {*a*_2_} in the system, and calculated the intergroup distance *d*({*x⃗*}) between their centers of mass as follows:

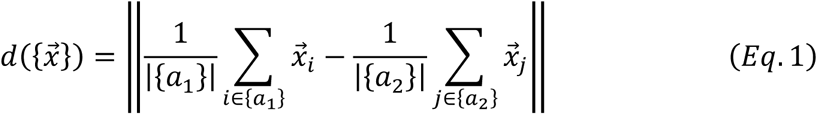

where *x⃗*_*i*_ represents the position vector of atom *i*, and |{·}| denotes the number of atoms in the group. Given a target intergroup distance *d*_*θ*_, we defined the error function as:

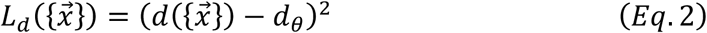

In the restraint-guided inference framework, we minimized *L*_*d*_({*x⃗*}) at each step of the reverse diffusion process to update the atomic coordinates. We applied this distance restraint method to two cases: 1) modeling changes in interdomain interactions within a protein, and 2) modeling changes in interactions between a protein and a ligand. The distance-restraint method was implemented in Boltz-2^22^, and all calculations were performed on NVIDIA RTX 4090 or RTX 5090 GPUs.

### Protein conformational change sampling and comparative evaluation

We first sampled conformational changes in proteins known to undergo transitions between two distinct structural states: Open and Closed conformations. Within each protein, we defined two regions, A and B, that remain rigid during the conformational transition, and used the center-of-mass distance as the reaction coordinate (Table 1). We denote the intergroup distances measured in the experimentally determined Open and Closed structures as *d*_open_ and *d*_closed_, respectively. We defined the sampling range by setting an upper bound *d*_max_ and a lower bound *d*_min_ for the reaction coordinate *d* and divided this range into four equal intervals, yielding five equally spaced sampling points including the endpoints. In this study, we set *d*_max_ = *d*_open_ and *d*_min_ = *d*_closed_ based on the crystal structures obtained from the PDB (Table 1). Structure prediction was performed using each *d* value as *d*_*θ*_ in Eq. 2. For each *d*_*θ*_ value, we generated nine structures using different random seeds, yielding a total of 45 samples per target. The Multiple Sequence Alignment (MSA) used as input was the complete, unsubsampled MSA generated by MMSeqs2^23^ via colabfold_search in LocalColabFold^24^.

**Table 1.** Experimental conditions for the distance-restraint method.

| Target | $d_{\text{closed}}$ (Å) | $d_{\text{open}}$ (Å) | Selection1 (residues) | Selection2 (residues) | Total MSA |
| --- | --- | --- | --- | --- | --- |
| QBP | 25.0 | 29.1 | 4-83,<br>185-223 | 89-179 | 9,604 |
| MBP | 27.4 | 30.3 | 0-108,<br>263-308 | 113-257,<br>315-369 | 4,770 |
| ADK | 19.7 | 35.9 | 29-58 | 121-158 | 16,866 |
| MDM2/p53 | 13.0 | 70.0 | Chain A (MDM2) | Chain B (p53) | 1,389/59 |

To benchmark the distance-restraint method, we compared it with other approaches for predicting alternative conformational states. Modifying the input MSA in AlphaFold2 has been used to predict different structural states. Among these approaches, MSA subsampling is a representative technique that generates structural diversity by partially reducing coevolutionary information^12^. AlphaFold3 and Boltz-2 are also capable of conformational change sampling through MSA subsampling^25,26^. Another approach is BioEmu, a deep learning model specifically designed and trained to sample protein conformational changes^18^. We performed structure predictions under seven different conditions for comparison: AlphaFold3, Boltz-2, and AlphaFold2 with either subsampled or complete MSA as input, and BioEmu with its default settings. For conditions using MSA subsampling, we set the number of sequences in the MSA used for prediction (MSA depth) to 16, 32, 64, 128, or 256, resulting in five MSA depths. Following the protocol in a previous study^12^, the number of recycles was set to 1. To minimize sampling bias, we created three sets of subsampled MSAs by randomly extracting sequences from the original MSA with different random seeds. In addition, structure predictions were performed with three different random seeds, yielding a total of 45 samples per target. For conditions without MSA subsampling, we used the complete MSA and performed 45 structure predictions with default options using different random seeds.

We employed multiple metrics to evaluate the predicted structures. To quantify structural similarity to reference structures, we calculated Template Modeling (TM) scores against the experimentally determined Open and Closed structures using US-align^27^. To assess prediction confidence, we evaluated confidence scores such as pLDDT (predicted Local Distance Difference Test) and pTM (predicted Template Modeling), provided by structure prediction methods including AlphaFold2, AlphaFold3, and Boltz-2. Furthermore, to assess the stereochemical validity, we analyzed structural quality metrics, including Angle RMSD, Bond RMSD, and Clashscore, using MolProbity^28^.

As model systems, we used three proteins with well-characterized conformational changes: glutamine-binding protein (QBP; Open structure: PDB ID 1GGG^29^, Closed structure: PDB ID 1WDN^30^), maltose-binding protein (MBP; Open structure: PDB ID 1OMP^31^, Closed structure: PDB ID 1ANF^32^), and adenylate kinase (ADK; Open structure: PDB ID 4AKE^33^, Closed structure: PDB ID 1AKE^34^). These proteins exhibit large-scale domain motions upon ligand binding and are well suited for evaluating the performance of conformational change sampling methods.

### Structural sampling along ligand dissociation pathways

We next applied the distance-restraint method to sample structures along the dissociation pathway of a protein-ligand complex. As a model system, we selected the MDM2-p53 complex^35^ (PDB ID: 1YCQ). In this system, we defined the reaction coordinate *d* as the distance between the respective centers of mass of MDM2 and the ligand (transactivation domain of p53). The lower bound of the reaction coordinate (*d*_min_) was set to 13 Å, corresponding to the distance in the experimental structure, while the upper bound (*d*_max_) was set to 70 Å (Table 1). The range between *d*_min_ and *d*_max_ was divided into nine equal intervals, yielding 10 equally spaced sampling points including the endpoints. Structure prediction was performed using each *d* value as *d*_*θ*_ in Eq. 2. We generated structures with five different random seeds for each *d* value, yielding a total of 50 structures. The generated structures were evaluated using confidence metrics, including pLDDT and interface pTM (ipTM), as well as stereochemistry-based quality metrics calculated with MolProbity.

### MD simulations for enhanced conformational sampling

To enable more detailed conformational sampling and subsequent thermodynamic analysis, we integrated deep learning-based structure prediction with physics-based MD simulations. We performed MD simulations under constant-temperature, constant-pressure conditions, using each structure generated by the distance-restraint method as the initial configuration. The N- and C-termini of proteins were capped with acetyl (ACE) and *N*-methylamide (NME) groups, respectively. Systems were prepared using the tleap module of AmberTools25^36^, employing the ff14SB^37^ force field for proteins. Solvation was performed using the TIP3P water model^38^ with a NaCl concentration of 0.15 M. The prepared systems were converted to GROMACS format using ParmEd^39^, and all MD simulations were conducted using GROMACS 2025.1^40^. Equilibration was first performed with positional restraints applied to the backbone atoms of protein and peptide ligand. Following equilibration, a production run was conducted for each initial structure. During both equilibration and production runs, temperature was controlled using the v-rescale method^41^ and pressure using the c-rescale method^42^.

### Markov state model construction and free energy estimation

Structural features, including the TM-score, were extracted from the MD trajectories, and MSMs^9^ were constructed to quantitatively evaluate free energy changes along the reaction coordinate using the Deeptime 0.4.5 package^43^. The MSM construction proceeded by extracting features from trajectories, clustering structures into discrete microstates, and computing the transition probability matrix *T*(*τ*) at lag time *τ*:

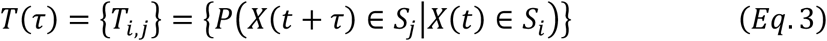

where *X* represents the feature vector extracted from simulation trajectories, and each element *T*_*i,j*_ denotes the transition probability from the discretized state *S*_*i*_ to *S*_j_. The Markovian property of the constructed models was validated by plotting the implied timescales (ITS) as a function of lag time τ and confirming the presence of a plateau:

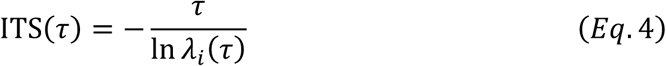

where *λ*_*i*_(*τ*) denotes the *i*-th eigenvalue of *T*(*τ*). Free energy changes along the reaction coordinate were evaluated using the stationary distribution obtained from the MSM. The potential of mean force (PMF) was calculated using the stationary distribution *π*, obtained by solving the eigenvalue problem with an eigenvalue of unity:

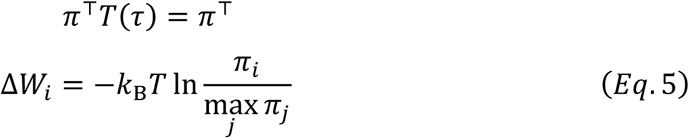

TM-scores against Open and Closed reference structures were calculated by US-align^27^ and used to construct two-dimensional free energy landscapes weighted by the MSM equilibrium distribution.

### US/MBAR analysis for free energy calculation

To obtain detailed free energy profiles along the reaction coordinate, we performed US simulations^21^ combined with the MBAR method^10^. Initial structures were generated using the distance-restraint method with finer distance intervals to ensure sufficient overlap between neighboring umbrella windows. For protein conformational changes, structures were generated at 0.1 Å intervals from *d*_min_ to *d*_max_ along the interdomain distance coordinate using a single random seed. For the protein-ligand dissociation process, structures were generated at 0.5 Å intervals along the inter-chain center-of-mass distance.

During US, a harmonic biasing potential was applied to restrain the reaction coordinate *d* to the target value *d*_*θi*_ for each window *i*:

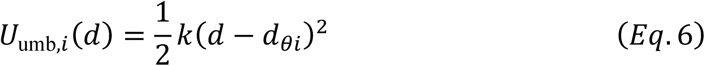

The force constants *k* were set to 4000, 6000, 2000, and 500 kJ mol⁻¹ nm⁻² for QBP, MBP, ADK, and the MDM2-p53 complex, respectively. These values were chosen to ensure sufficient overlap between neighboring windows while maintaining adequate sampling within each window. The US simulations were performed using the standard pull code functionality in GROMACS.

Free energy profiles were reconstructed using the MBAR method^10^, which provides statistically optimal estimates by solving the following self-consistent equation for the dimensionless free energies *f*_*i*_:

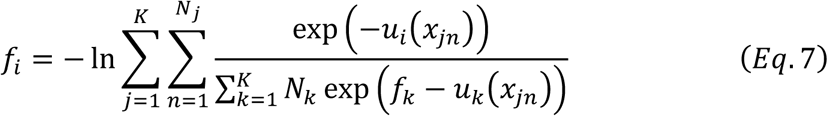

where *u*_*i*_(*x*_*jn*_) is the reduced potential of configuration *x*_*jn*_ sampled from window *j* and evaluated under potential *i*, *K* is the total number of windows, and *N_j_* is the number of samples from window *j*. Since the system potential energy cancels out between windows, only the biasing potential *U*_umb_ needs to be considered in the analysis^44^. To reduce computational cost and avoid bias from correlated samples, statistically independent samples were extracted from each trajectory based on autocorrelation analysis prior to the MBAR calculation. All MBAR analyses were performed using pymbar 4.0.3.

## Results

### Evaluation of protein conformational sampling strategies

To evaluate the conformational sampling capability of our distance-restraint method, we compared it with seven alternative conditions: AlphaFold3, Boltz-2, and AlphaFold2, each tested with either subsampled or complete MSA as input, as well as BioEmu with default settings. For each structure predicted under these conditions, we calculated TM-scores against both the Open and Closed reference structures of the target proteins and visualized these scores as two-dimensional plots to systematically assess the conformational sampling performance (Figure 2).

**Figure 2.**
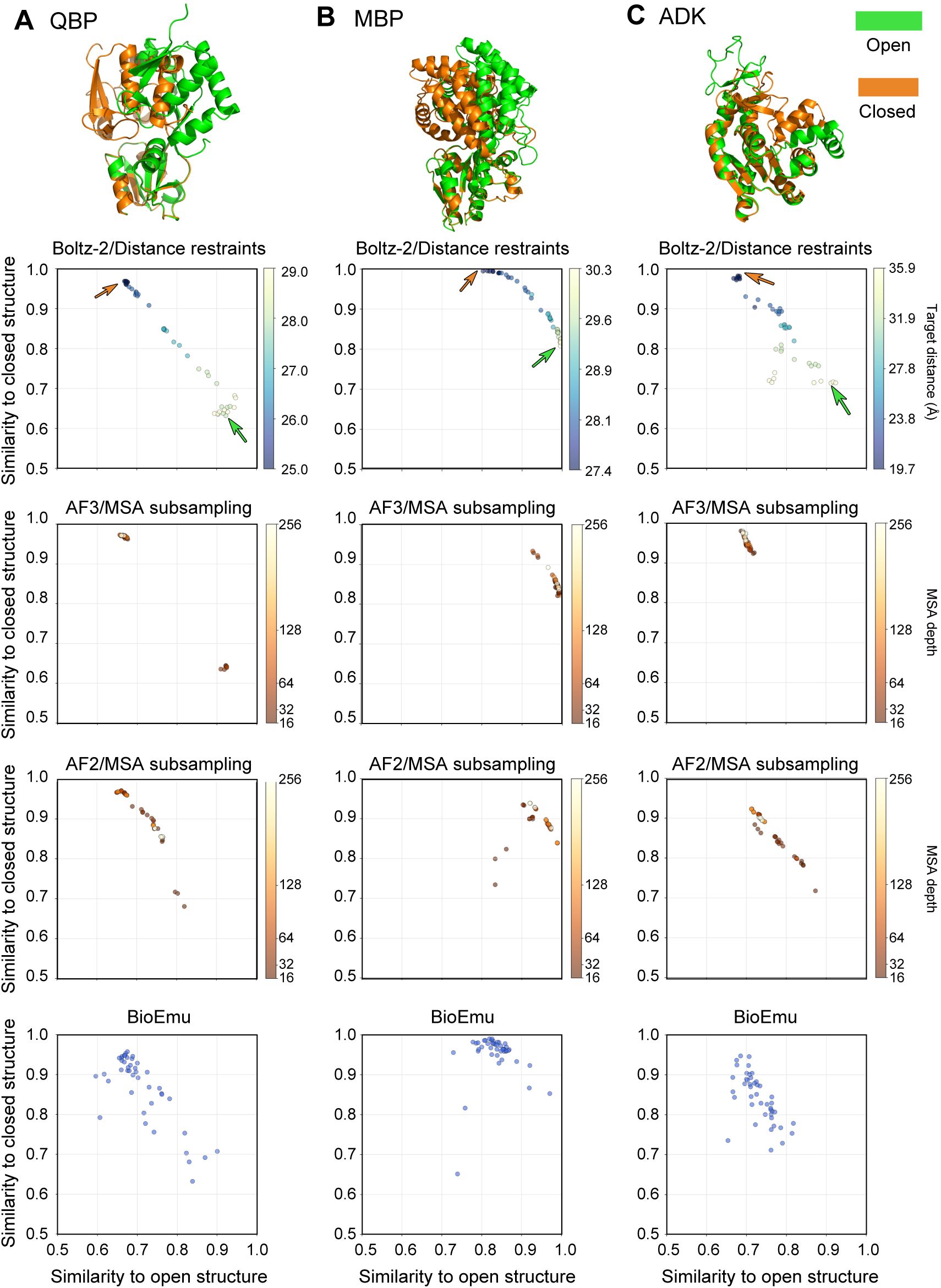
Conformational sampling of three model proteins by different methods. (A) QBP, (B) MBP, and (C) ADK. Top panels show superimposed crystal structures of Open (green) and Closed (orange) conformations. Scatter plots display TM-scores of predicted structures against the Open (x-axis) and Closed (y-axis) reference structures, showing the region where both TM-scores are ≥ 0.5. For the distance-restraint method implemented in Boltz-2, points are colored by target restraint distance (Å), with arrows indicating the positions of the Open and Closed crystal structures. For AlphaFold3 and AlphaFold2 with subsampled MSA, points are colored by MSA depth (number of sequences).

Our distance-restraint method successfully sampled conformational transitions from Closed to Open states along the restraint distance *d* in all three test cases: QBP, MBP, and ADK (Figure 2). Notably, the method achieved uniform sampling across the conformational space, including intermediate states. This demonstrates that explicit control of the reaction coordinate enables systematic sampling along the conformational transition pathway. In contrast, AlphaFold3, Boltz-2, and AlphaFold2 with subsampled MSA failed to achieve complete coverage of the conformational transition pathway (Figure 2). These methods tended to generate structures biased toward either the Open or Closed state, with particularly limited coverage observed for ADK and QBP. Relatively uniform sampling was achieved for MBP, suggesting that the effectiveness of MSA subsampling is protein-dependent. These results highlight the difficulty of appropriately controlling coevolutionary signals that contribute to the stability of Open and Closed states when using MSA subsampling approaches. Consequently, achieving desired conformational states through MSA adjustment must rely on empirical trial and error. Interestingly, standard AlphaFold3 using a complete MSA without modification exhibited some degree of conformational diversity for MBP simply by sampling the latent space of the diffusion model (Figure S1). Similary, standard Boltz-2 with a complete MSA captured part of the conformational transition in ADK, albeit with a biased distribution (Figure S1). These observations indicate that the diffusion models used in AlphaFold3 and Boltz-2 can inherently generate some structural diversity through sampling of their latent spaces, as recently reported^25^. However, in all cases, the sampling efficiencies were inferior to those achieved with the distance-restraint method.

BioEmu is trained not only on static experimental structures but also on trajectories obtained from MD simulations with thermodynamic weighting^18^, and is therefore expected to generate protein structures that reflect thermodynamic ensembles. Analysis of structures generated by BioEmu revealed broader sampling of conformational space than the other conventional methods. However, the sampled region remained more limited than that achieved by the distance-restraint method. Furthermore, BioEmu generated some structures in regions that deviated substantially from those sampled by other methods. Visual inspection revealed that many of these outliers exhibited physically unrealistic conformations for proteins at room temperature, including significantly stretched bond lengths (Figure S2).

To assess the quality of predicted structures, we analyzed pLDDT and pTM scores, which are learning-based confidence metrics calculated by default in AlphaFold2, AlphaFold3, and Boltz-2. Although some variation was observed among methods, all approaches—including the distance-restraint method, MSA subsampling, and complete MSA conditions—produced structures with confidence scores within acceptable ranges (*i.e.*, pLDDT > 80, pTM > 0.7), indicating that structural reliability was maintained irrespective of the prediction strategy employed (Figure S3). Note that BioEmu does not provide pLDDT or pTM scores, so these metrics could not be evaluated for this method. We further evaluated the generated structures using MolProbity^28^, which calculates stereochemistry-based quality metrics. For the distance-restraint method, all three test cases (QBP, MBP, and ADK) showed favorable MolProbity scores, and individual metrics were also within acceptable ranges (Figure S4). This result indicates that the application of distance restraints does not compromise the stereochemical validity of protein structures. Similarly, favorable stereochemical metrics (i.e., Angle RMSD, Bond RMSD, and Clashscore) were obtained for all methods except BioEmu, irrespective of MSA subsampling (Figure S4). In contrast, the stereochemical metrics for BioEmu-generated structures were notably worse than those for other methods (Figure S4). This can be attributed to the sampling of physically unrealistic conformations, which resulted in severely compromised stereochemistry that in turn degraded the overall quality metrics.

We next examined the applicable range of the distance-restraint method using QBP as an example, by investigating the effects of imposing restraint distances beyond the appropriate range determined from the Open and Closed experimental structures. When the restraint range was extended to span from 20 Å to 40 Å for *d*_min_ and *d*_max_, we observed an overall decrease in pLDDT scores at distance outside the appropriate range, accompanied by deterioration of MolProbity quality indicators including Angle RMSD, Bond RMSD, and Clashscore (Figure S5). These results suggest that even for systems where experimental structures are unavailable, monitoring these quality metrics could enable empirical determination of biologically valid restraint ranges.

### Free energy landscapes of protein conformational changes

Structures predicted by AlphaFold2, AlphaFold3, and Boltz-2, with or without distance restraints, are sampled from model distributions trained on experimental biomolecular structures. Although these distributions are not identical to the Boltzmann distribution of the target system under ambient conditions, regions of high likelihood within the learned distributions are expected to correspond reasonably well to thermodynamically favorable conformational states. To approximate the true Boltzmann distribution and enable quantitative thermodynamic analysis, we performed MD simulations starting from the predicted structures to sample their local conformational neighborhoods. The choice of simulation strategy depends on the specific application: conventional MD (cMD) simulations combined with MSM analysis^9^ are well suited for validating whether structures generated by distance restraints reside in thermodynamically stable regions without introducing additional biases, whereas US/MBAR^10^ provides an effective approach for detailed sampling of transition regions and precise free energy quantification.

For MSM-based analysis, We sampled multiple structures using the distance-restraint method and performed 10 ns cMD simulations starting from each structure for QBP, MBP, and ADK. Trajectory data from all simulations were combined, and MSMs were constructed using TM-scores against the Open and Closed crystal structures as feature variables. The number of clusters and the lag time were set to 100 and 5 ns, respectively, and the Markov property was verified using implied timescales (Figure S6). Two-dimensional free energy landscapes were then derived from the stationary distributions of the MSMs (Figure 3). The initial structures sampled by our method corresponded to regions within 0–3 kcal/mol on the free energy landscape, demonstrating that the distance-restraint method successfully samples thermodynamically stable conformations (Figure 3). Examination of these landscapes indicated that the Open state represents the stable ground state in the apo form, whereas the Closed state is energetically less favorable (Figure 3).

**Figure 3.**
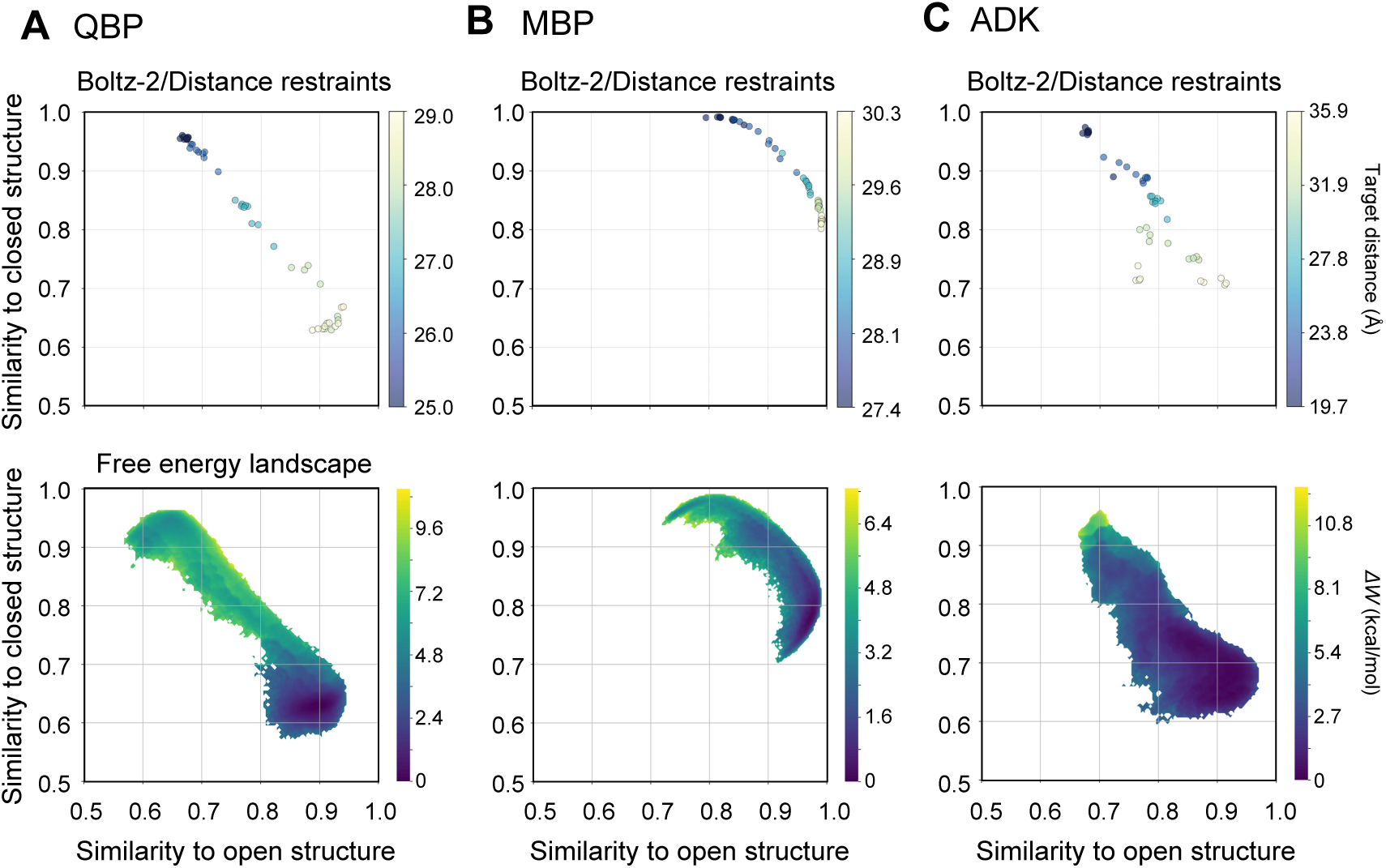
Integration of the distance-restraint method and MD simulations for free energy landscape construction using Markov state models. (A) QBP, (B) MBP, and (C) ADK. Upper panels: structures after MD equilibration, plotted by TM-scores against open (x-axis) and closed (y-axis) reference structures. Points are colored by target restraint distance (Å). Brown and green arrows indicate the closed and open states, respectively. Lower panels: two-dimensional free energy landscapes derived from MSM equilibrium distributions. Color scale indicates relative free energy *ΔW* (kcal/mol).

To obtain quantitative free energy changes along the conformational transition pathway, we performed US/MBAR analysis using the interdomain distance as the reaction coordinate, consistent with the target distance *d* used in the distance-restraint method. The resulting one-dimensional free energy profiles revealed that the free energy difference from the Open ground state to the Closed state (defined at the interdomain distance of the Closed crystal structure, *d*_closed_) was 6.24 ± 0.25 kcal/mol for QBP, 5.81 ± 0.29 kcal/mol for MBP, and 4.64 ± 0.10 kcal/mol for ADK (Figure 4). Since both the distance restraint method and subsequent MD simulations were performed on apo proteins, the greater stability of the Open state is consistent with expectation. The US/MBAR analysis was also consistent with the protein-specific behavior observed in the MSM landscapes: although QBP exhibited a flat shoulder in the free energy profile near the Closed state, none of the three proteins showed a distinct local minimum corresponding to a stable Closed conformation in the ligand-free state (Figure 4).

**Figure 4.**
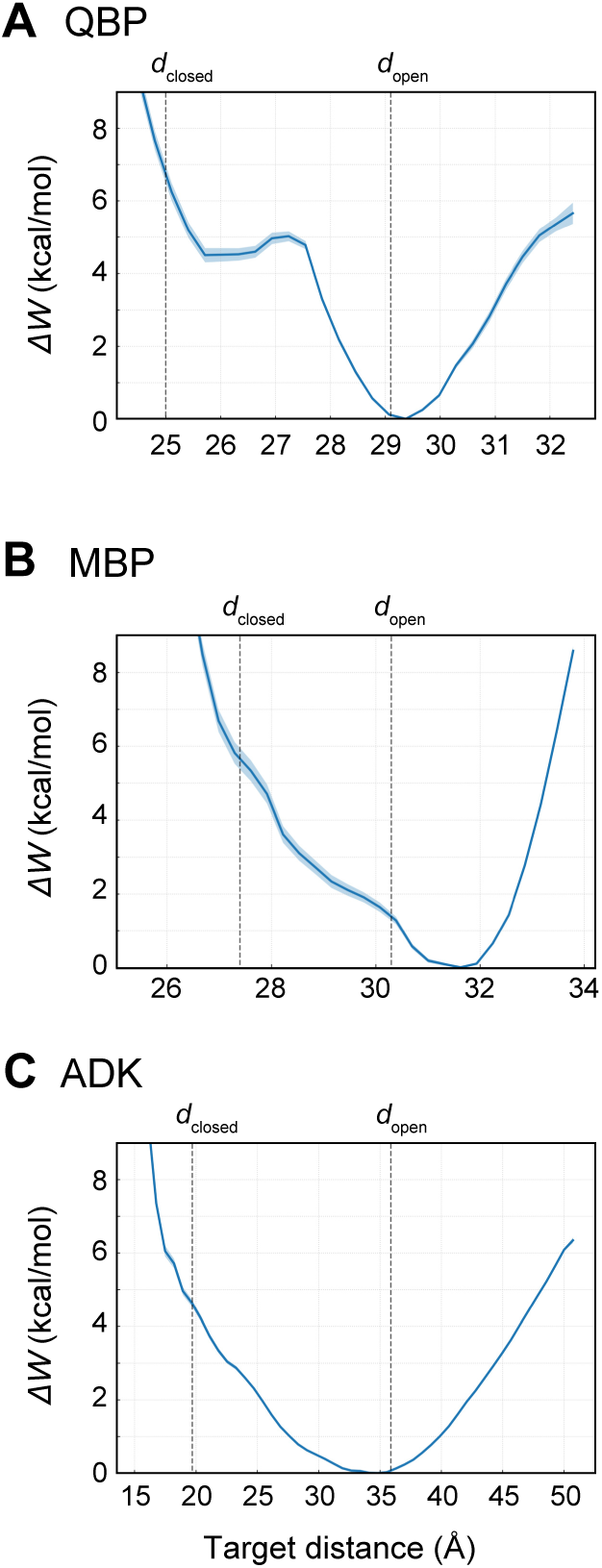
Free energy profiles along the interdomain distance calculated by US/MBAR. Free energy profiles (*ΔW*) as a function of the interdomain center-of-mass distance for (A) QBP, (B) MBP, and (C) ADK. Vertical dashed lines indicate the *d* value corresponding to the Closed and Open states, *d*_closed_ and *d*_open_, respectively. Shaded regions represent statistical uncertainties.

### Sampling and analysis of protein-ligand dissociation

The distance-restraint method developed in this study can control not only intramolecular domain motions within proteins but also the relative positioning between proteins and other biomolecules. As an application of this capability, we sampled the structural pathway of protein-ligand dissociation. To validate the effectiveness of the method, we selected the MDM2-p53 system, in which the ligand is a peptide with substantial molecular weight. Existing structure prediction models typically output bound-state structures, reflecting the predominance of stable complex structures in their training data. In contrast, our method systematically varied the center-of-mass distance between protein and ligand as a restraint condition, successfully generating an ensemble of complex structures at arbitrary distances, including intermediate states (Figure 5). To assess whether the generated structures exhibited any abnormalities such as distorted protein conformations or severe steric clashes between protein and ligand, we performed MolProbity analysis (Figure S8). The results showed favorable MolProbity scores in all cases, confirming that the structures generated under distance restraints are stereochemically valid. We further investigated how prediction confidence scores changed with restraint distance. The pLDDT score remained in a favorable range throughout, indicating that the structures were not corrupted (Figure S7). However, interface-related scores such as ipTM decreased as the restraint distance increased. This trend is attributable to the fact that the underlying model was primarily trained on bound complex structures; consequently, dissociated states represent out-of-distribution configurations.

**Figure 5.**
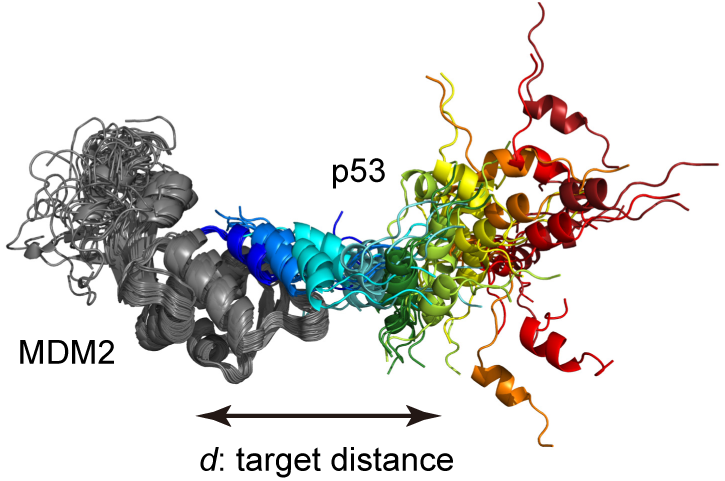
Structural ensemble of the MDM2-p53 complex sampled by the distance-restraint method. Overlay of predicted structures for the MDM2-p53 system with varying target distances d between the centers of mass of MDM2 and the p53 peptide. MDM2 is shown in gray, and the p53 peptide conformations are colored from blue to red according to increasing target distance (13–70 Å). The double-headed arrow indicates the reaction coordinate *d*.

We next combined the structural ensemble efficiently sampled by the distance-restraint method with physics-based MD simulations to quantitatively evaluate the free energy landscape associated with ligand binding. For the MDM2-p53 system, as with the conformational change cases described above, we used multiple structures sampled along the reaction coordinate by the distance-restraint method as initial configurations. US simulations were subsequently performed using these structures as starting points, with each window running for 50 ns. Trajectory data from all simulations were analyzed using the MBAR method, and a one-dimensional free energy profile was generated using the center-of-mass distance between MDM2 and p53 as the reaction coordinate. The resulting free energy profile showed a plateau where the energy converged to a constant value at center-of-mass distances exceeding approximately 30 Å (Figure 6). This plateau corresponds to complete dissociation where interactions between MDM2 and the p53 peptide are lost. The potential of mean force (*ΔW*) from the bound to unbound state calculated from the US/MBAR analysis of the MDM2-p53 system was 11.6 ± 0.42 kcal/mol, which is consistent with values reported in previous studies^45^. These results demonstrate that the proposed approach, combining pathway exploration through the distance-restraint method with subsequent MD simulation and free energy analysis, provides an effective strategy for thermodynamic quantification of dynamic biomolecular processes.

**Figure 6.**
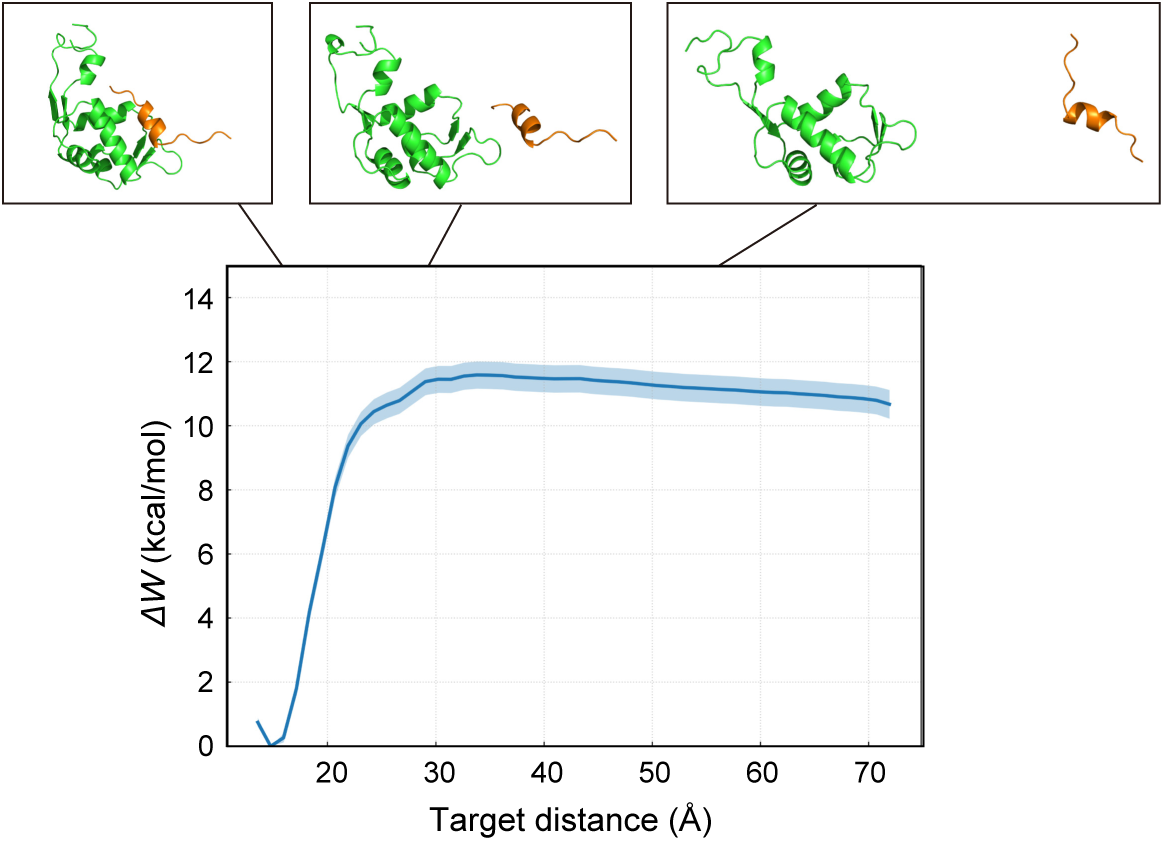
Free energy profile of MDM2-p53 dissociation. One-dimensional free energy profile as a function of the center-of-mass distance between MDM2 and the p53 peptide. The solid line represents the potential of mean force, with shading indicating the statistical uncertainty. Insets show representative structures at the indicated distances along the dissociation pathway, with MDM2 shown in green and the p53 peptide in orange.

## Discussion

In this study, we implemented the distance-restraint method in Boltz-2, and successfully reproduced protein conformational changes and ligand dissociation/association processes without retraining the model. By integrating this deep learning-based generative sampling approach with MD simulations, we achieved broader sampling of the Boltzmann distribution and enabled more quantitative evaluation of biomolecular conformational dynamics.

The distance restraints applied during the reverse diffusion process play a role analogous to US in MD simulations, using a harmonic bias potential along a predefined reaction coordinate to enhance sampling at specified coordinate values. Recent studies have begun to explore physics-inspired modifications to the generative trajectories of diffusion models for biomolecular structure prediction^22^, such as metadynamics-like approaches that progressively discourage previously sampled conformations to promote structural diversity^46^. While these metadynamics-inspired methods can generate diverse structures, they inherently lack precise control over which conformational states are sampled. In contrast, our US-like approach enables targeted generation at arbitrary points along a chosen reaction coordinate. This controllability also provides clear advantages over MSA-based structure prediction methods. Compared with the MSA manipulation approaches, our method achieves more systematic sampling of conformational states and remains applicable to protein-ligand complexes involving small molecules, where no coevolutionary signal is available. Moreover, MSA-based methods that reduce coevolutionary signals to induce conformational diversity may inadvertently perturb regions that do not undergo conformational change, potentially distorting predicted structures in these stable domains (Figure S9). Our method preserves the complete MSA and coevolutionary information, thereby avoiding this issue.

Despite these advantages, the proposed method has several limitations. Conformational changes that cannot be well characterized by intergroup distances remain challenging for our method. For example, the activation of G protein-coupled receptors (GPCRs) involves complex and concerted movements of multiple transmembrane helices, making it impractical to define a low-dimensional reaction coordinate using intergroup distances. For such complex conformational changes, conventional MSA manipulation methods, despite their inability to uniformly sample conformational pathways, may still provide a valid alternative. Additionally, the distance-restraint method requires prior knowledge of which domains undergo movement, whereas MSA-based approaches may explore unknown conformational changes through reduction of coevolutionary signals.

Extending this method to multi-dimensional reaction coordinates represents an important direction for future development. For instance, capturing protein conformational changes accompanying ligand dissociation would require simultaneous distance restraints on both the ligand-protein distance and the protein interdomain distance. A key challenge in this extension is developing strategies to efficiently explore only meaningful regions corresponding to low free energy states, as the search space grows exponentially with increasing number of reaction coordinates.

Beyond MSM-based analysis, our method may facilitate other free energy calculation approaches such as the string method^47,48^, which require initial structures distributed along the reaction coordinate. These structures have conventionally been generated using steered MD or targeted MD simulations, which are computationally demanding and prone to hysteresis effects; our approach offers a rapid alternative without such artifacts. Another potential application lies in structure-based virtual screening, where cofolding approaches using AlphaFold3-like models are increasingly explored as alternatives to conventional docking methods. A known limitation of the cofolding methods is the difficulty in controlling which binding pocket the ligand occupies. While current implementations provide options to specify binding pockets^22^, our distance-restraint method could offer more robust control by explicitly guiding ligands toward specific sites, particularly for targeting cryptic or allosteric binding sites.

## Data and Software Availability

The materials supporting this article, including the source code, will be available on GitHub (https://github.com/cddlab/boltz_restr).

## Author contributions

T. H., Y. M., and R.I. contributed to the design of the work, and T.H. performed computational experiments and wrote the manuscripts, and R.I. supervised this study.

## Acknowledgements

This research was partially supported by JST BOOST under Grant Number JPMJBS2430 for T.H. and by the Research Support Project for Life Science and Drug Discovery (Basis for Supporting Innovative Drug Discovery and Life Science Research (BINDS)) from AMED under Grant Numbers JP25ama121027 for Y.M. and JP25ama121012 for R.I. This work was also supported by JSPS KAKENHI Grant-in-Aid for Transformative Research Areas (A) under Grant Numbers 25H01570 and 25H02250 for R.I. Additional support was provided by the Medical Research Center Initiative for High Depth Omics at the Institute of Science Tokyo, Nanken-Kyoten at the Institute of Science Tokyo 2025, and the Multilayered Stress Diseases project (JPMXP1323015483) at the Institute of Science Tokyo.

---

**Figure S1.**
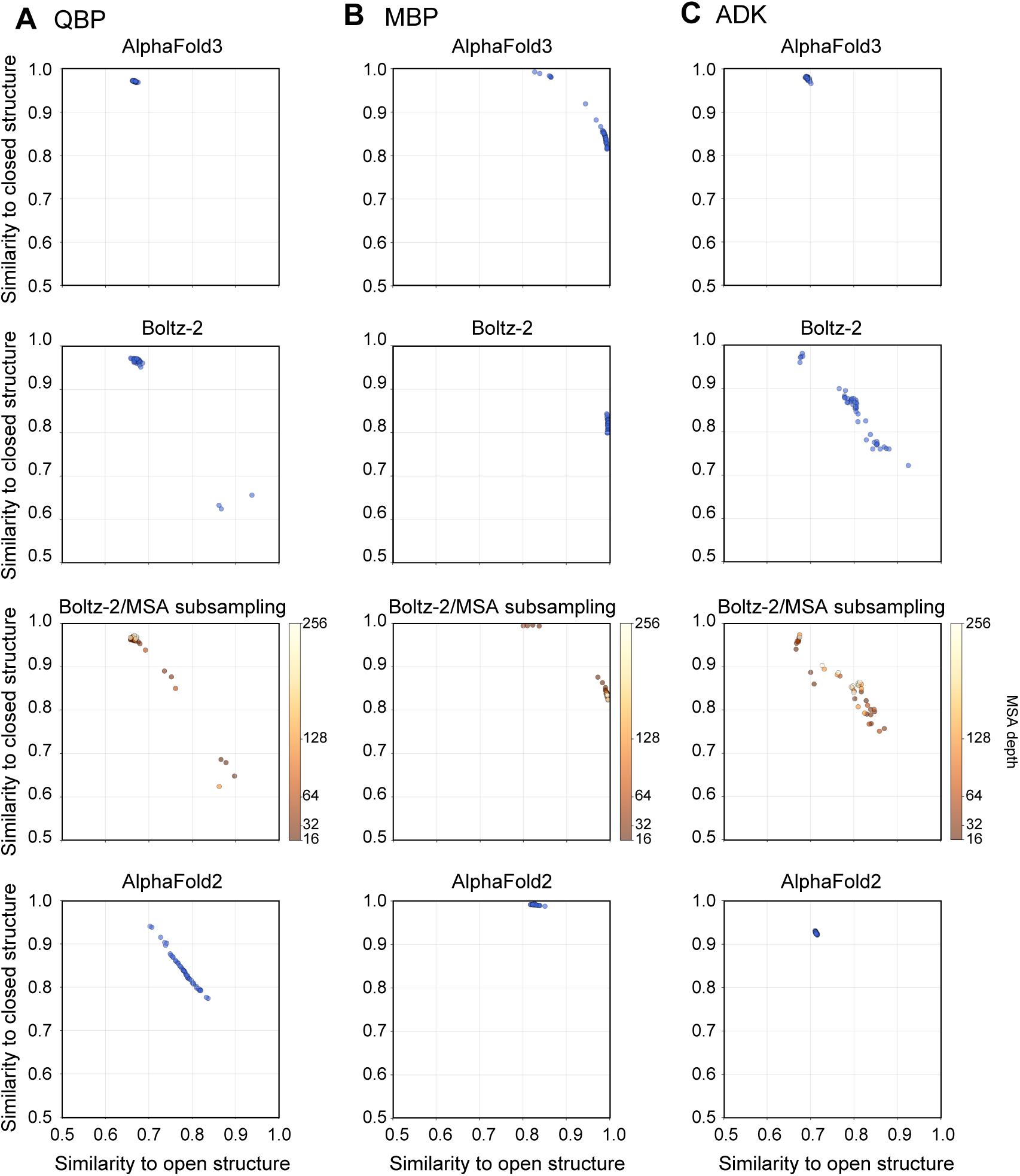

**Figure S2.**
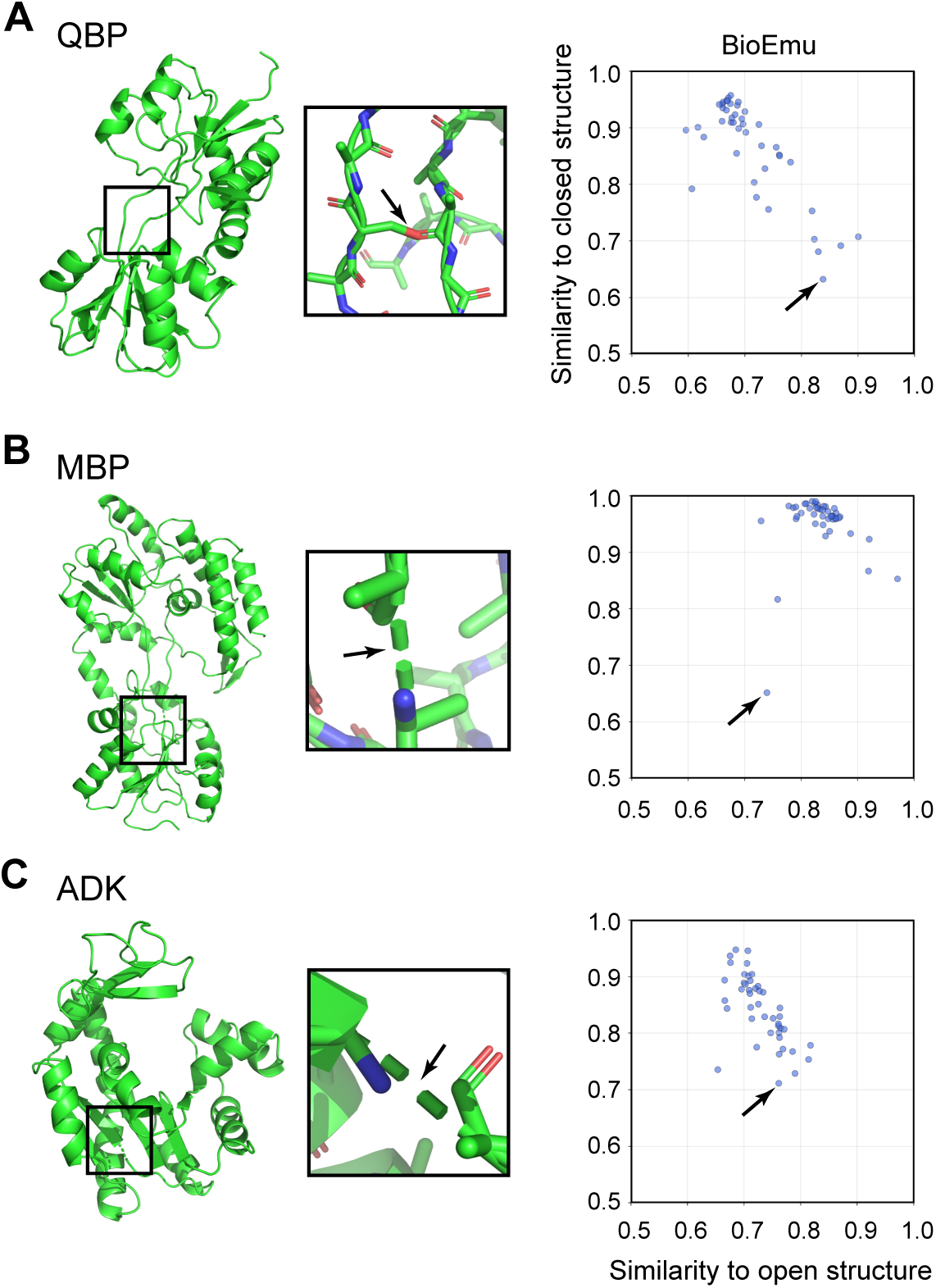

**Figure S3.**
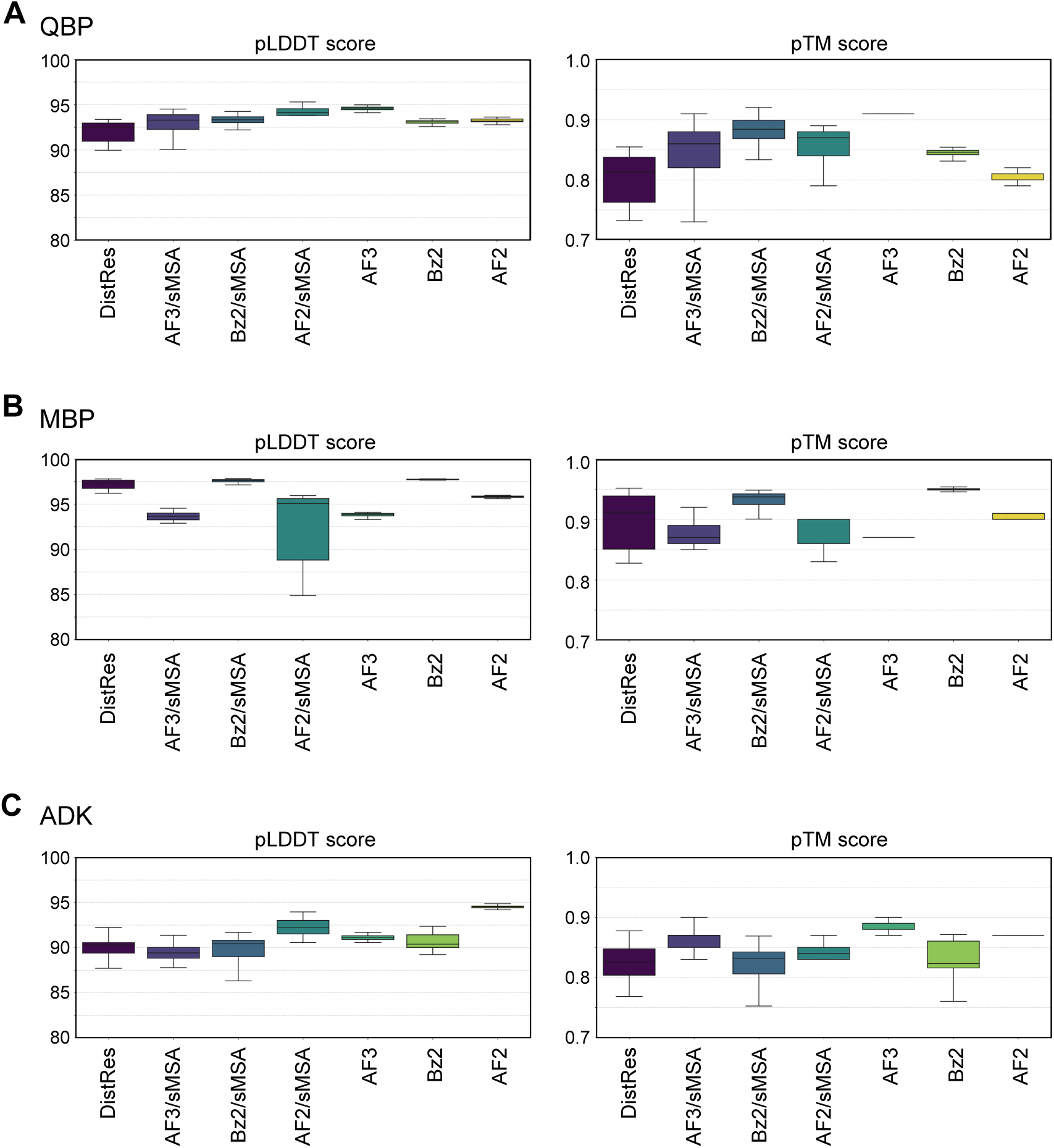

**Figure S4.**
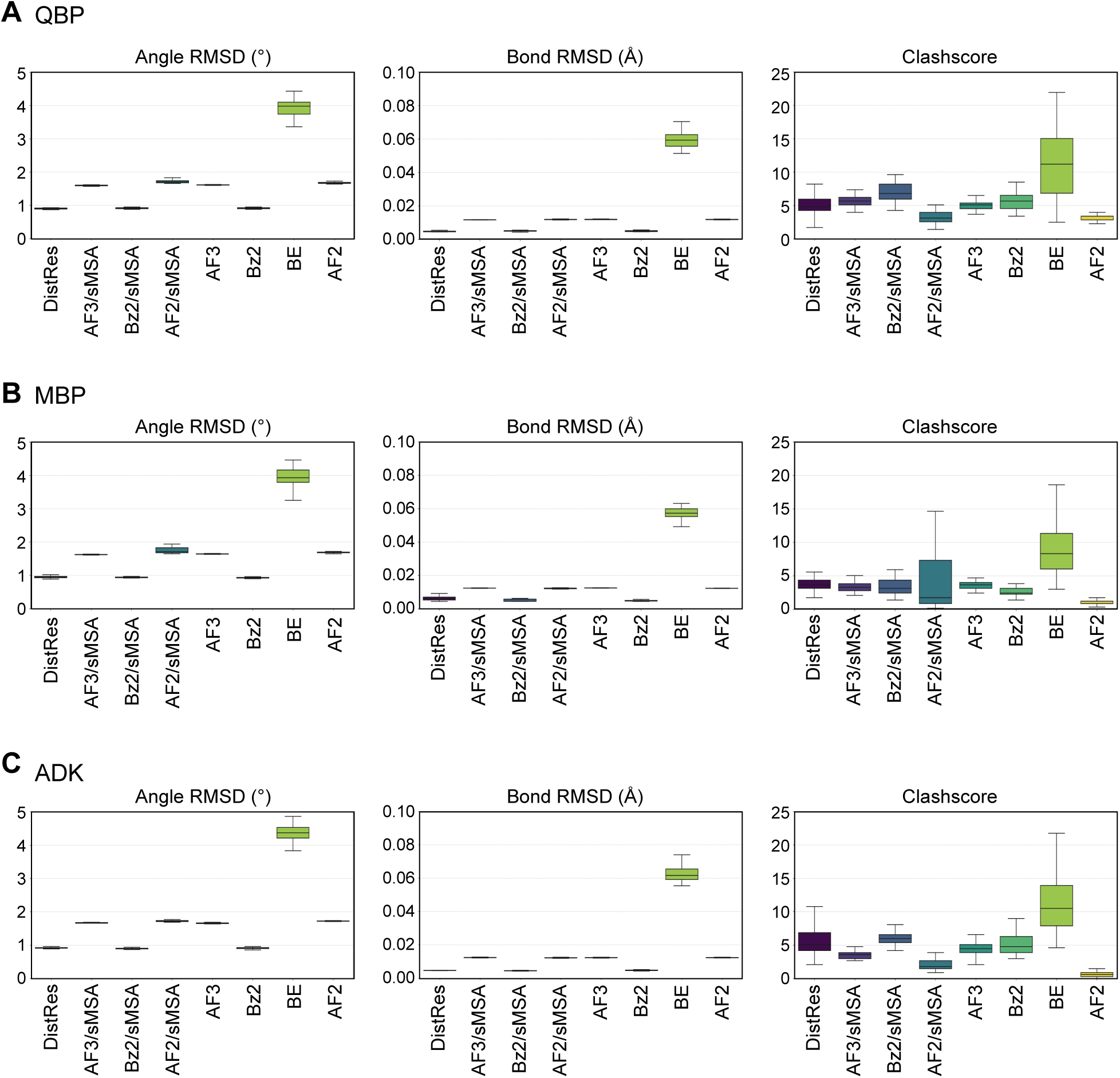

**Figure S5.**
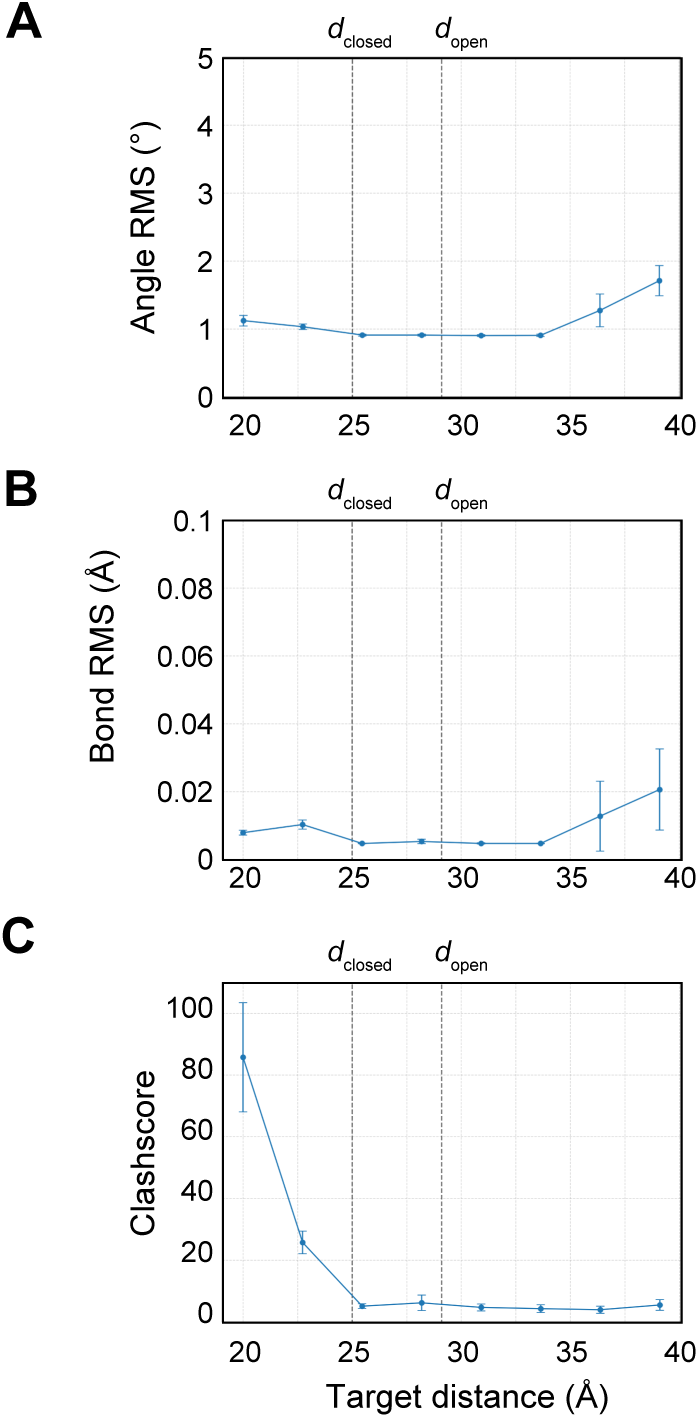

**Figure S6.**
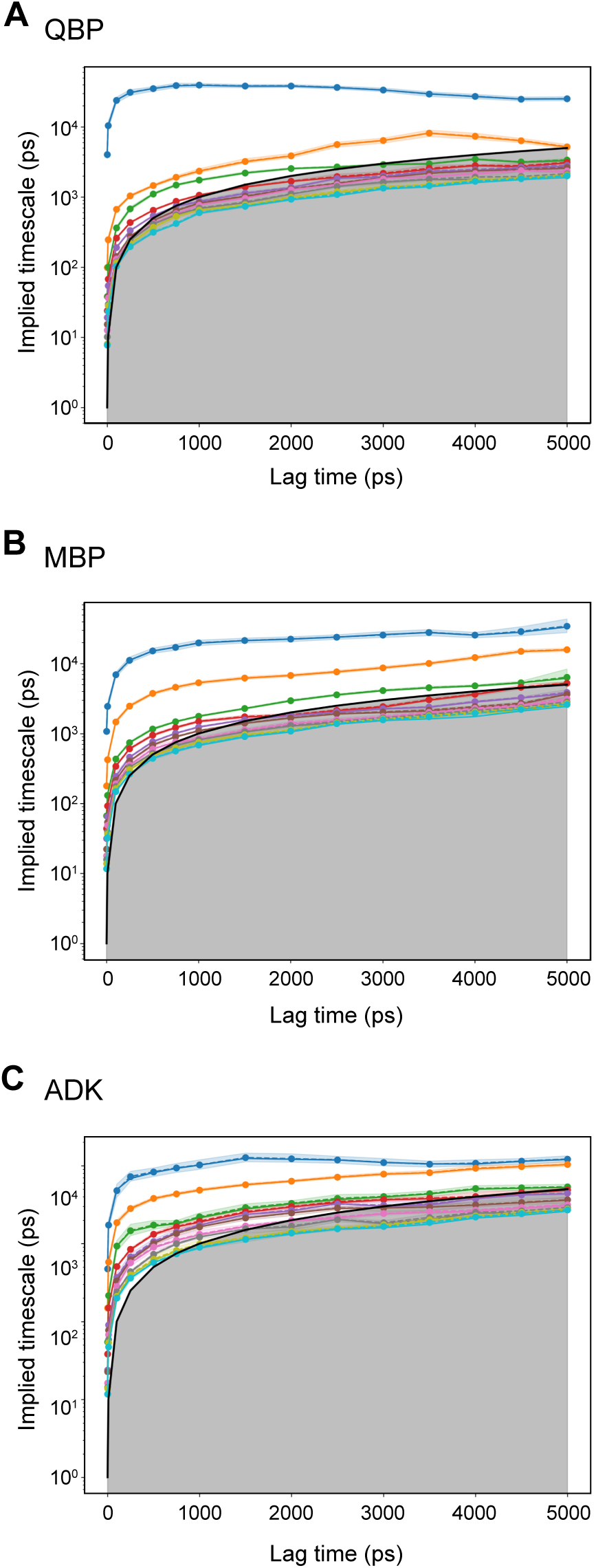

**Figure S7.**
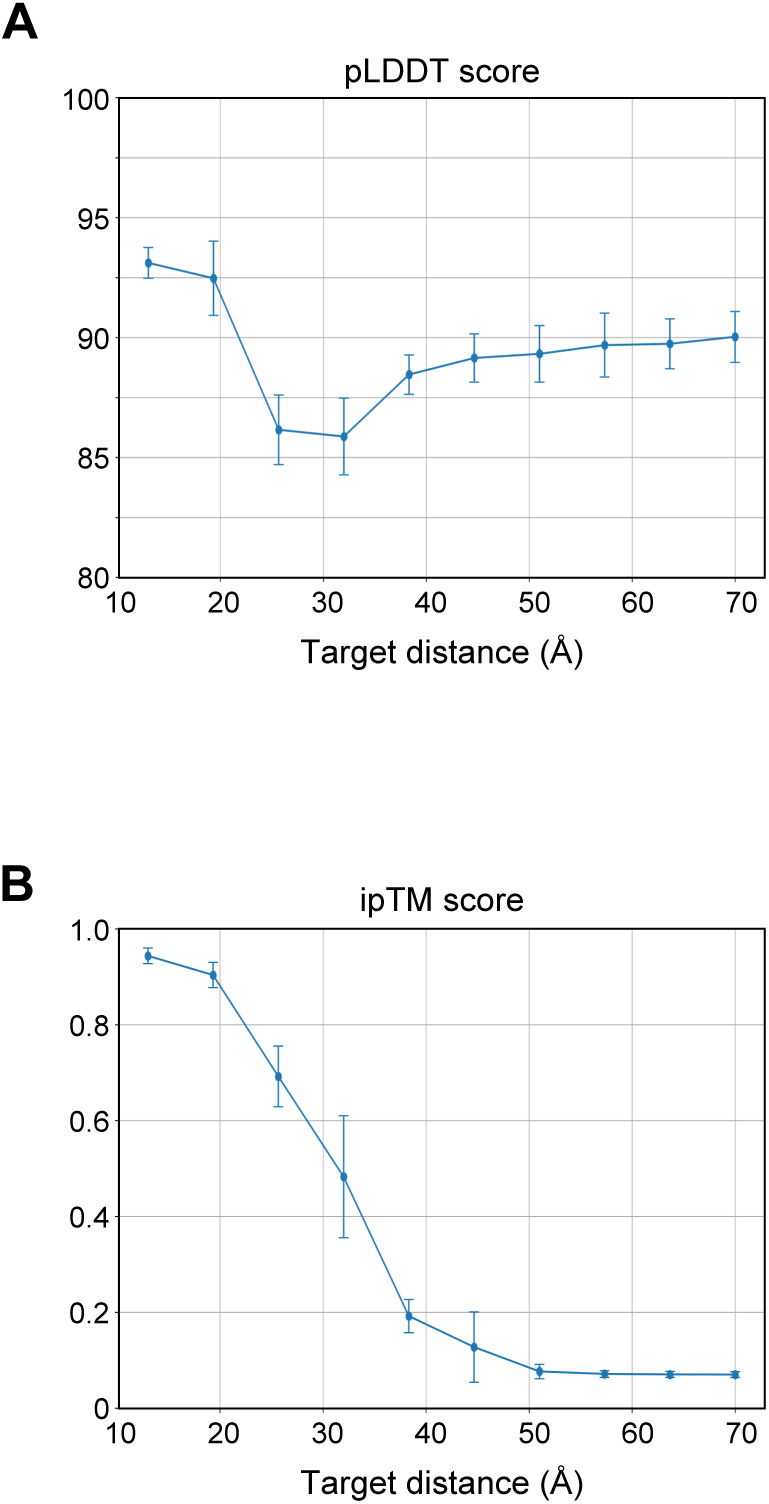

**Figure S8.**
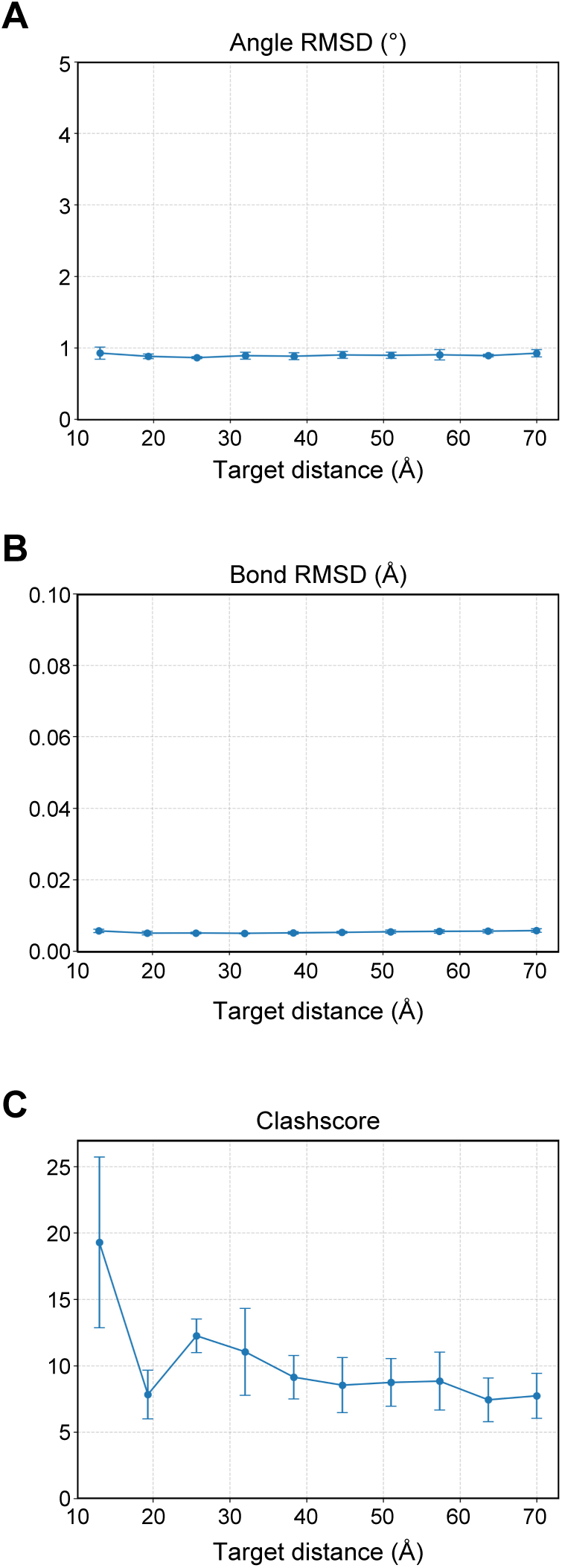

**Figure S9.**
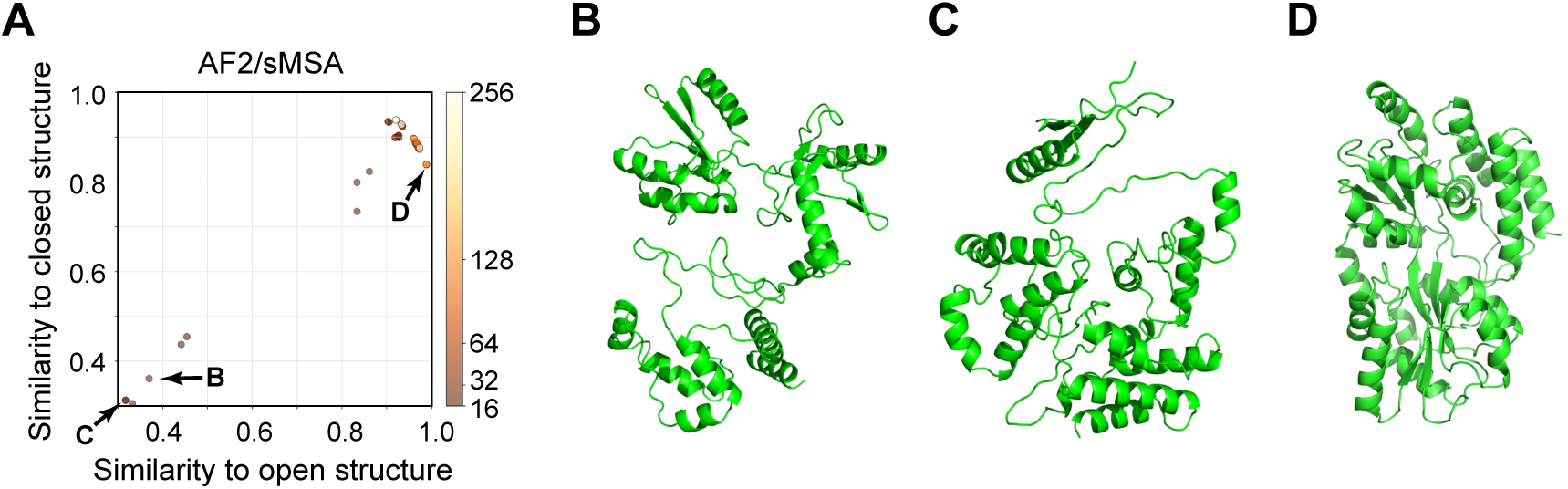

