## Supplementary material for "Distance-Restraint-Guided Diffusion Models for Sampling Protein Conformational Changes and Ligand Dissociation Pathways": suppl

### Supplementary Figure Legends

#### Figure S1

Conformational sampling by conventional structure prediction methods.

Scatter plots showing TM-scores of predicted structures against Open (x-axis) and Closed (y-axis) reference structures. Columns correspond to QBP (A), MBP (B), and ADK (C), while rows show results from AlphaFold3, AlphaFold2, Boltz-2, and, Boltz-2 with MSA subsampling (top to bottom). For Boltz-2 with MSA subsampling, point colors indicate MSA depth as shown in the color bar.

#### Figure S2

Outlier structures generated by BioEmu.

(A) QBP, (B) MBP, and (C) ADK. Left panels show representative outlier structures generated by BioEmu. Insets display magnified views of the boxed regions; arrows highlight specific structural defects such as severely stretched bond lengths. Scatter plots show the distribution of sampled structures in TM-score space, with similarity to the open reference structure on the x-axis and similarity to the closed reference structure on the y-axis.

#### Figure S3

Learning-based confidence scores for predicted structures across conformational sampling methods.

Box plots showing pLDDT scores (left) and pTM scores (right) for structures predicted by each method. (A) QBP, (B) MBP, (C) ADK. DistRes, Boltz-2 with distance restraints; AF3/sMSA, AlphaFold3 with MSA subsampling; Bz2/sMSA, Boltz-2 with MSA subsampling; AF2/sMSA, AlphaFold2 with MSA subsampling; AF3, AlphaFold3 with complete MSA; Bz2, Boltz-2 with complete MSA; AF2, AlphaFold2 with complete MSA. Boxes indicate interquartile ranges with median lines; whiskers extend to 1.5× interquartile range.

**Figure S4**

Stereochemical quality assessment of predicted structures for protein conformational sampling.

Box plots showing MolProbity quality metrics for structures generated by different prediction methods. (A) QBP, (B) MBP, and (C) ADK. Three metrics are shown for each protein: Angle RMSD (°), Bond RMSD (Å), and Clashscore. Methods compared are: DistRes, Boltz-2 with distance restraints; AF3/sMSA, AlphaFold3 with MSA subsampling; Bz2/sMSA, Boltz-2 with MSA subsampling; AF2/sMSA, AlphaFold2 with MSA subsampling; AF3, AlphaFold3; Bz2, Boltz-2; BE, BioEmu; AF2, AlphaFold2. Boxes indicate interquartile ranges with median lines; whiskers extend to 1.5× interquartile range.

**Figure S5**

Stereochemical quality metrics as a function of restraint distance in protein conformational sampling.

(A) Angle RMSD, (B) Bond RMSD, and (C) Clashscore for QBP plotted against target restraint distance. Vertical dashed lines indicate $d_{\text{closed}}$ and $d_{\text{open}}$ corresponding to the experimentally determined Closed and Open structures, respectively. Data points represent mean values with error bars indicating standard deviations across nine independent predictions at each target distance.

### Figure S6

Implied timescale analysis for Markov state model validation.

Implied timescales (ITS) as a function of lag time for MSMs constructed from MD simulations of (A) QBP, (B) MBP, and (C) ADK. The ten slowest relaxation modes are shown. The gray shaded region indicates the range below the lag time, where ITS values are not meaningful.

### Figure S7

Learning-based confidence scores for distance-restrained ligand dissociation sampling.

(A) pLDDT scores and (B) ipTM scores for MDM2-p53 complex structures plotted against target restraint distance d, defined as the center-of-mass distance between protein and ligand. Data points represent mean values, and error bars indicate standard deviations across five independent predictions at each restraint distance.

### Figure S8

Stereochemical quality assessment of predicted structures for distance-restrained ligand dissociation sampling.

MolProbity quality indicators for MDM2-p53 complex structures generated at varying restraint distances. (A) Angle RMSD, (B) Bond RMSD, and (C) Clashscore plotted against target distance between protein and ligand centers of mass. Data points represent mean values, and error bars indicate standard deviations across structures generated at each restraint distance.

### Figure S9

Structural distortion in AlphaFold2 predictions with reduced MSA depth.

(A) TM-scores of structures predicted by AlphaFold2 with MSA subsampling (AF2/sMSA) against the Open (x-axis) and Closed (y-axis) reference structures of MBP. Points are colored by MSA depth. Arrows indicate samples corresponding to structures shown in panels B–D. (B, C) Representative structures with low TM-scores to both Open and Closed references, exhibiting significant structural distortion. (D) The predicted structure with high TM-score to the Open reference, maintaining proper fold integrity.
